# Quantitative Nanohistology of aging dermal collagen

**DOI:** 10.1101/2023.03.02.530877

**Authors:** Sophia Huang, Adam Strange, Anna Maeva, Samera Siddiqui, Phillipe Bastien, Sebastian Aguayo, Mina Vaez, Hubert Montagu-Pollock, Marion Ghibaudo, Anne Potter, Herve Pageon, Laurent Bozec

**Affiliations:** Faculty of Dentistry, University of Toronto, Toronto, Ontario, Canada; Eastman Dental Institute, University College London, London, United Kingdom; L’Oréal Research & Innovation, Aulnay-sous-Bois, France; School of Dentistry, Faculty of Medicine, Pontificia Universidad Catolica de Chile, Chile; Institute for Biological and Medical Engineering, Schools of Engineering, Medicine and Biological Sciences, Pontificia Universidad Católica de Chile; Physics Department, Lancaster University, Lancaster’ United Kingdom

**Keywords:** Collagen, Ageing, Biomarkers, Atomic Force Microscopy, indentation, Principal Component Analysis, nano-Histology

## Abstract

While the external signs of skin aging have been well-defined throughout history, much less is known about aging within the ultrastructure of our skin. Our skin, the largest organ in our body, is structured by collagen through fibrils or large sheets. With the increased use of nanometrology tools in histology, it is now possible to explore how the aging process affects collagen at its most fundamental level, the collagen fibril. Here, we show how atomic force microscopy-based quantitative nanohistology can differentiate skin from different age groups and anatomical sites. Following the definition of specific collagen biomarkers at the nanoscale, we used a segmentation approach to quantify the prevalence of 4 structural biomarkers over a dataset of 42,000 images (30 donors) complemented by extensive nanomechanical analyses (30,000 indentation curves) on histological sections. Our results demonstrate that specific age-related collagen fingerprints could be found when comparing the % prevalence of each marker between the papillary and reticular dermis. A case of abnormal biological aging validated our markers and nanohistology approach. This first extensive study focusing on defining signs of dermal aging at the nanoscale proves we are all unique to our dermal collagen ultrastructure.

## INTRODUCTION

The skin is a complex and multi-layered connective tissue that defines the external appearance of all individuals. Specific pathologies or the unavoidable aging process directly impact this appearance. Genetic influences and internal factors such as hormones or metabolic substances1 govern the intrinsic aging of the skin. It illustrates the naturally occurring skin modifications with age, leading to wrinkles and skin dryness. The wrinkle formation in human skin has been associated with marked decreases in skin elasticity^2^. Extrinsic skin aging arrives earlier and is due, for example, to exposure to sunlight, pollution, or lifestyle choices such as a lack of balanced nutrition ^3^. More generally, in aging skin, cell replacement is continuously declining, the barrier function and mechanical protection are compromised, wound healing and immune responses are delayed, thermoregulation is impaired, and sweat and sebum productions are decreased.

The aging of the dermis is a complex process with several changes occurring to both collagen and non-collagenous components of the dermal ECM. Both collagen and elastin undergo enzymatic and non-enzymatic crosslinks in the ECM as a function of aging^4,5^. Lysyl oxidase (LOX) is the primary enzyme that crosslinks collagen and maintains collagen alignment and fibril structure^6^. However, the proportion of LOX-derived crosslinks reduces with age ^7^, while the proportion of non-enzymatic (glycation) crosslinks increases with age. Accumulating non-enzymatic glycation products increases collagen crosslinking in the skin with age. Glucosepane^8^ is the most common AGE crosslink (between lysine and arginine) resulting from the interaction of the collagen molecule with glucose, the highest concentrated sugar found in vivo ^9^. Another characteristic of aging in the dermis is related to an elevation in MMP-1 activity, which is responsible for dermal collagen fibril fragmentation in aged human skin^10,11^. Evidence suggests that MMP-1-mediated cumulative collagen damage significantly contributes to the phenotype of aged human skin.

While the effects of aging of the skin at the macroscale are well-reported, these effects have been sparsely investigated on the collagen fibril organization in the dermis. An early study investigated how the nanoscale mechanical properties of the extracellular matrix regulate dermal fibroblast function ^12^. This study was the first to systematically explore mechanical and structural dermis properties (human) as a function of age (26 to 55). Alongside presenting the first ultra-topography images of native skin by Atomic Force Microscopy (AFM), they found that the dermis elasticity ranged from 0.1 to 10kPa in hydrated sections. These results were further explored by Ahmed et al. ^13^, who proposed the first phenotypic markers for collagen aging in the reticular dermis, also measured by AFM. In addition, Ahmed explored the relationship between advanced glycation products and the elasticity of individual collagen fibrils within the reticular dermis of young and older individuals. Thus, despite being one of the most imaged proteins by AFM, collagen remains largely uncharacterized at the fibrils scale within the dermis, especially as a function of aging. Here, we present the first large study (30 donors) in which we explore the use of AFM-based quantitative nanohistology to quantify the biophysical properties of dermal collagen at the nanoscale and how the variations in these properties can be used to reveal the skin’s biological age.

## MATERIALS AND METHODS

### Human skin histological sections

In this study, cryo-preserved historical skin samples (30 Caucasian female donors, 18 to 75 years, cosmetic surgery procedures) collected under informed consent were analyzed as part of this explorative study. All the samples were anonymized, and the donors’ age, biological sex, and ethnicity were made available for this study. The sample cohort was split into three groups according to both the donors’ age and anatomical site: <u>B</u>reast <u>Y</u>oung <u>S</u>kin (BYS, N=11), <u>B</u>reast <u>O</u>ld <u>S</u>kin (BOS, N=9), and <u>C</u>heek <u>O</u>ld <u>S</u>kin (COS, N=10). Selected human skin samples were fixed in neutral formalin and then embedded in paraffin or cryo-embedded OCT block before being sectioned (5um). The sections were then stained using Hematoxylin and Eosin Stain (HES), Orcein, and Sirius Red (SR). For quantitative nanohistology, four unstained cryo-sections (8-10 µm thickness) were physisorbed directly on individual glass slides.

### Histological Imaging

A Leica (Wetzlar, Germany) light microscope (LM) was used for histology imaging of the stained sections. This microscope was equipped with two crossed-light polarizers (90°) to allow for polarization (darkfield) LM and with an 8-megapixel digital camera (EOS Rebel 100, Canon). Complete histological sections were digitized using the auto-stitch function in Image-Pro Plus software (Meyer Instr. Inc, Houston, USA).

### Quantitative Nanohistological Imaging

Topological images (10×10μm^2^) of the tissue were acquired by Atomic Force Microscopes (AFM Nanowizard I & III, Bruker-JPK, Berlin – Germany) operated in contact mode (MSNL cantilevers, Bruker, Santa Barbara) and in ambient conditions directly on the unstained sections. To avoid bias in selecting the area to be imaged, we used a random walk approach to land the probe in each dermal layer by using the AFM sample holders’ x-y translation screws to move the sample (within each dermal layer). Following their acquisition, all AFM images were plane-fitted before being segmented into 1×1 μm^2^ images using Image J, creating a final topology dataset consisting of 42,000 (1×1 μm^2^) images.

### Quantitative Nanomechanical analysis

The mechanical properties of collagen fibrils of the histological sections were acquired by a Nanowizard I AFM (Bruker-JPK, Berlin – Germany) operated in the force-distance mode in ambient conditions. For these measurements, RFESPA cantilevers (Bruker, Santa Barbara) with a spring constant k= 3 N/m were employed. All indentations (1Hz, max load 350nN) were carried out exclusively on collagen fibrils exhibiting a clear and identifiable D-banding periodicity (min 500 individual collagen fibrils indented per dermal layer). The mechanical dataset consisted of 30,000 indentations for the 30 samples. All force-distance curves were processed by the JPK Data Processing Software v.5.1.8 using the Sneddon/Hertz model ^14^. The resultant distribution of Elastic (Young’s) Moduli was plotted as histograms with a fixed bin size (B=500MPa) to calculate the median Elastic (Young’s) Moduli for the respective dermal layers.

### Data Management & Statistical Analyses

All data were processed using Origin Pro (Origin Lab Corporation, Northampton, USA). In this study, each histological block was treated as an individual variable, and data obtained from serial histological sections from the same block were pooled together. Four calibrated (using 100 AFM 1×1 μm^2^ images) analysts were used to quantify the prevalence of the four structural biomarkers in each 42,000 1×1 μm^2^ image. The Young’s modulus of fibrils was compared using a two-sided hypothesis Mann-Whitney U test with a significance level α=0.001. In addition, 2D Principal component analyses^15^ (PCA) were performed on the dataset using Origin Pro (Origin Lab Corporation, Northampton, USA). The relative position of the variables on the PCA loading plot, with their coordinates equal to correlations with the first two principal components, was used to assess the level of discrimination between the variables. Finally, a Partial least Squares Discriminant Analysis (PLS-DA) was performed using SIMCA® 16.0 multivariate Data Analysis Software (Sartorius Stedim Data Analytics AB, Sweden) with the binary age class as the response variable.

## RESULTS & DISCUSSION

### Histology of an aging dermal section

Histological analysis is routinely used to assess structural changes in tissues, including skin, to prognose or diagnose pathologies such as aging. Figure 1 presents the histology of representative sections from our three groups. The young group (Breast Young Skin– BYS) represented by the photo-protected breast skin specimen presents distinct epidermis (Ep), papillary (PD), and reticular (RD) dermal layers (Fig. 1a). The dermo-epidermal junction is non-uniform and presents well-defined rete ridges (arrow). Structurally, the collagen in both dermal layers is very dense, and there is little evidence of elastin (Fig. 1b-i, iv, and vii). In the intrinsically aged skin (Breast Old Skin: BOS) specimen, one observes a thinning of the epidermis and papillary dermis (Fig. 1b-ii). The dermo-epidermal junction presents a flattening aspect resulting from the disappearance of the rete ridges. The dermis has an atrophic aspect with a loss of cells and extracellular matrix. As a result, the dermal collagen becomes sparser (Fig. 1b-v), and elastin accumulates in this intrinsically aged skin (Fig. 1b-viii).

**Figure 1.**
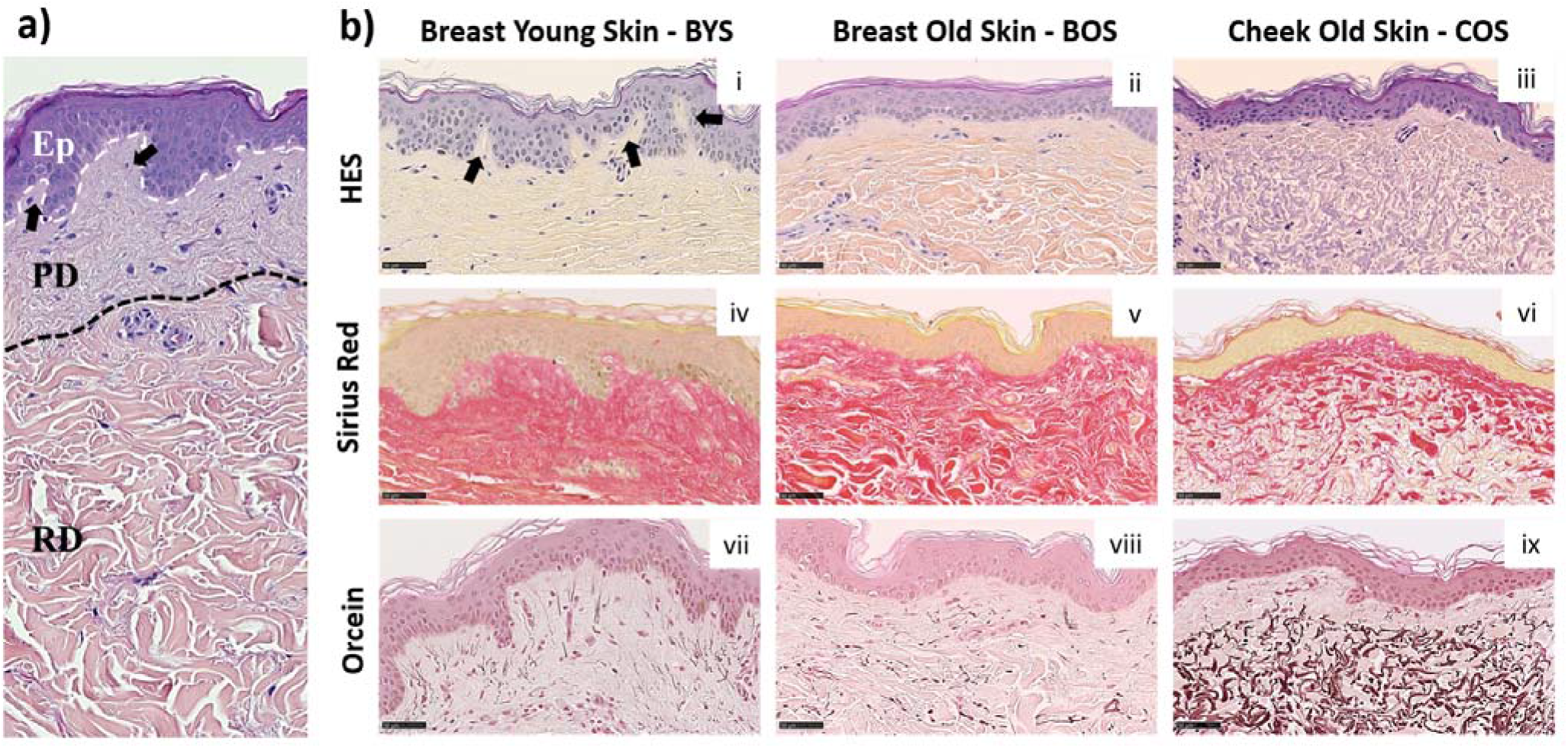
Histology of Normal Human Skin: (A) Normal human skin from a young donor (22 yo). The dotted lines separate the epidermis (Ep), papillary (PD), and reticular (RD) dermis. Histological colorations of human skin from breast young (i, iv, vii) and old (ii, v, viii) skin (respectively 22 and 74 yo) and cheek old (iii, vi, ix) skin (75 yo). Different types of colorations were performed: Hematoxylin Eosin Saffron – HES (i-iii), Sirius red (iv-vi), and Orcein (vii-ix). Scale Bar = 50 µm. Arrows highlight the rete ridges.

In contrast, the primary histological fingerprint for extrinsically aged skin (Cheek Old Skin: COS) is the abundance of abnormally structured elastin accumulated as elastotic material. This accumulation can be found in the reticular dermis, as shown in Fig 1b-ix. In addition, the extrinsically aged skin specimen presents a flattening of the dermis and a loss of rete ridges found for the intrinsically aged skin (Fig. 1b-iii). The ratio of elastin to collagen has also increased (Fig. 1b-vi, ix), suggesting that the collagen is being replaced by elastin, which explains the altered functional and biomechanical properties of this skin ^16^. The histological presentation of these groups: young photo-protected skin and intrinsically and extrinsically aged skin, are well known and are routinely used to define the pathological age of skin^17^.

### Defining nanoscale structural biomarkers for (type I) collagen

Type I collagen fibril topology is one of the most recognized protein structures due to its highly conserved annular-banding periodicity along the long axis of the fibrils, namely the D-banding periodicity ^18^. In our quantitative nanohistology approach ^19^, we have used AFM to image collagen structure on the skin groups’ histological sections. Figure 2 presents our approach to correlating AFM images site with a polarized image of SR-stained skin sections. We observe the presence of long fibrils with a defined D-banding periodicity in all the images recorded. Fibrils can appear as thick bundles, homogeneous sheets, or interwoven scaffolds. Our original skin study defined several potential structural biomarkers for aging in collagen ^13^. Four were found to be present across the three skin groups from a list of 7 candidates’ structural (or topological) collagen biomarkers, as presented in Figure 2b. Marker 1 is defined as the presence of interfibrillar gaps, suggesting a loosening of the dense collagen sheet structure, leading to the loss in fibril registration with one another (Fig. 2b-i). Marker 2 characterizes areas where the fibrillar collagen structure is not readily observable and could be associated with cell processes or other extracellular matrix components (Fig. 2b-ii). Marker 3 characterizes a well-defined collagen matrix, with the fibrillar D-banding apparent (Fig. 2b-iii). However, the collagen fibrils in those areas are disorganized and lack registration between them. In other studies, we found collagen fibrils morphology prevalent in the fibrotic area ^19^. Finally, Marker 4 characterizes a well-defined collagen matrix, with the fibrillar D-banding apparent and the fibrils aligned, forming dense collagen sheets (Fig. 2b-iv). The prevalence of each of these structural (or topological) collagen biomarkers in the papillary and reticular dermis by segmenting each 10×10 μm^2^ AFM image into a 100 sub-image 1×1 μm^2^ as presented in Figure 3a. Each sub-image was evaluated independently (1 chosen marker per 1×1 μm^2^ image). In addition to these structural biomarkers, we added the median value of the collagen fibrils Young’s modulus for both the papillary and reticular dermis, as presented in Figure 3b. We have already demonstrated the variation of Young’s modulus of collagen fibrils (dry) as a function of the aging process ^13,20^.

**Figure 2.**
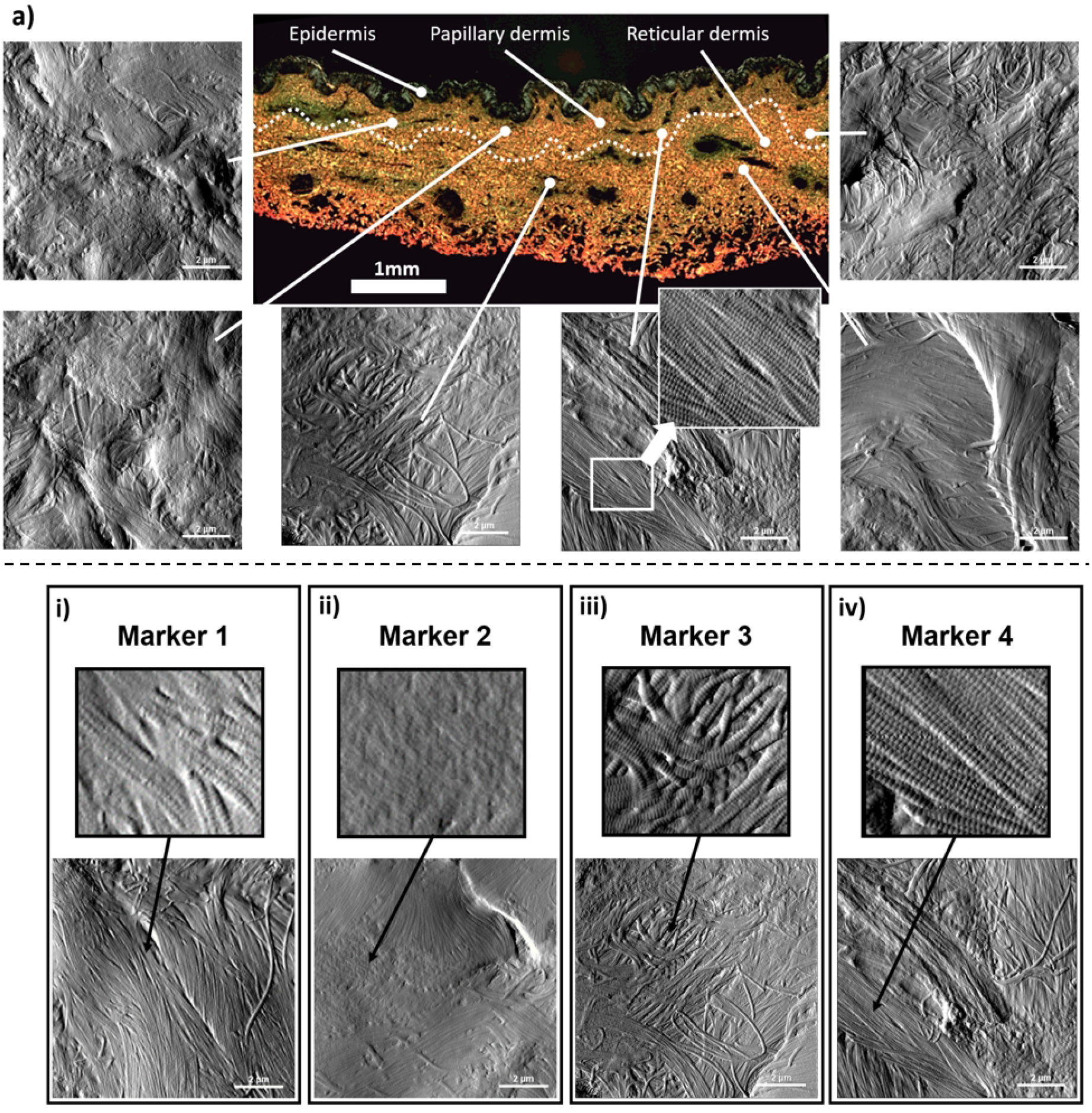
Quantitative Nanohistology of dermal collagen performed on histological sections: a) Representative localized AFM topological images (10 x10 μm^2^) obtained from both the papillary and reticular dermal layers obtained directly on a polarised Sirius Red Stain histological section. The boundary between papillary and reticular dermal layers, denoted as a dashed line, was empirically set 500μm below the epidermal junction. b) Structural nanoscale biomarkers for (type I) collagen and their presentation on AFM topological images: i) Marker 1 is defined as the presence of interfibrillar gaps (or holes); ii) Marker 2 characterizes areas where the fibrillar collagen structure is not readily observable and could be associated with cell processes or other extracellular matrix components; iii) Marker 3 characterizes a well-defined collagen matrix, with the fibrillar D-banding apparent on the collagen fibrils, but presenting a disorganized and lack registration between the collagen fibrils; iv) Marker 4 characterizes a well-defined collagen matrix, with the fibrillar D-banding apparent on the fibrils and forming a dense sheet of aligned collagen fibrils.

**Figure 3.**
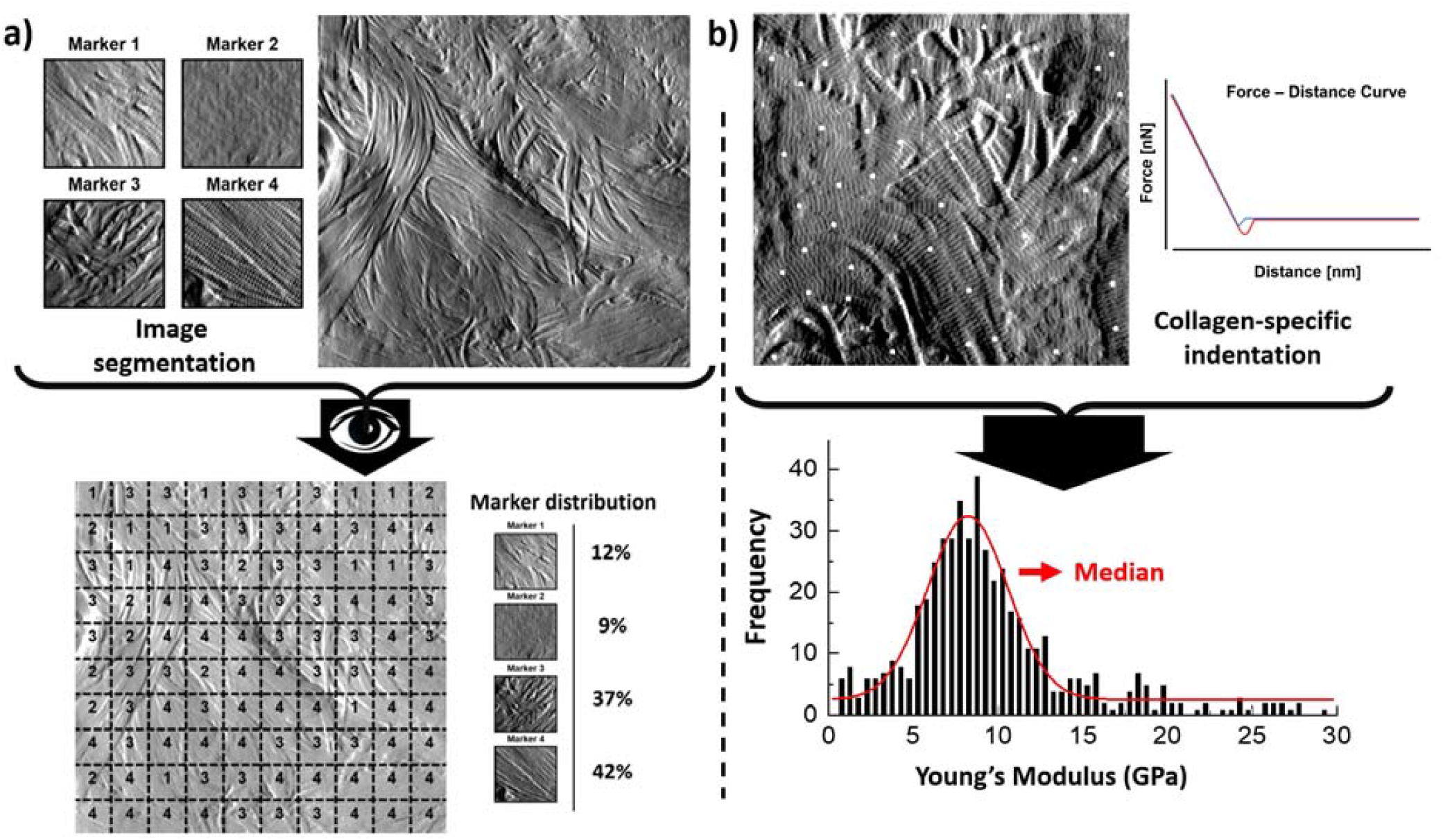
Quantitative Nanohistology image analysis and mechanical data analysis: **a**) AFM-based image visual segmentation approach presenting the four collagen morphological markers and a representative AFM topological image pre and post-visual segmentation, and the resultant marker distribution. The segmented AFM topological image is analyzed by overlaying a 10×10 macro pixel grid and assigning a unique marker in each of the grid areas; b) specific collagen fibrils AFM-based nanoindentation presenting localized indentation points performed on fibrils (white dots) to generate the frequency distribution of the Young’s Moduli which is in turn fitted to extract the median Young’s Moduli (compressive) of the fibrils indented in the AFM image.

### Deciphering the structure of collagen in dermal layers

Dermal collagen phenotype varies among individuals based on biological sex, ethnic origins, and lifestyles ^2, 1^. To analyze collagen fibril phenotype, one cannot assume that the properties of the dermal collagen for all the individual donors within a given group are similar; thus, markers cannot be represented through a common median value per dermal layer and donor group.

We explored whether the relative structural variations in the collagen matrices between the papillary and reticular dermis could be used to differentiate between the three groups. To do so, we referenced the variations in individual markers in the reticular dermis to those found for the papillary dermis and normalized these variations using Eq.1 in Figure 4-a. The results are plotted in Figure 4-a, presenting the %variation of each marker becoming more prevalent towards either the papillary or reticular dermis. Both markers 1 and 2 show an increased prevalence in the papillary dermis of the BYS group. While marker 2 does not readily inform us of the collagen phenotype, marker 1 suggests that the collagen matrix in the papillary dermis of young photo-protected skin exhibits structural loosening due to the increased presence of interfibrillar gaps (or holes) when compared to the reticular dermis. This finding would also suggest that the collagen matrix aging would occur in the papillary dermis before progressing to the reticular dermis. This outcome supports the recent finding by Lynch et al., who established that the mechanical property of the papillary dermis decreases before that of the reticular dermis with age ^21^. Their study suggested that with aging, the earliest microstructural and mechanical changes occur in the topmost layers of the dermis/skin and then propagate deeper, providing an opportunity for topical preventive treatments acting at the level of the papillary dermis. An earlier study focusing solely on phenotyping the reticular dermis collagen ^13^ suggested marker 3 is the hallmark of collagen aging. In our present study, marker 3 does not appear to be more prevalent in either dermal layer, regardless of age. This result does not contradict our previous study but implies that at the nanoscale, one cannot differentiate the two dermal layers solely based upon marker 3. Marker 4 shows an increased prevalence in the reticular dermis of the BOS group, suggesting that intrinsic aging promotes the formation of localized, well-aligned collagen fibril bundles. Using our nanohistology approach, we can assert that the reticular collagen is more aligned than the papillary collagen in the sparser region of dermal collagen. This increase in fibril alignment can be directly associated with interfibrillar crosslinks, such as those mediated by the formation of advanced glycation end-products ^22^. Finally, no clear trends exist in the % variation of any of the markers associated with the COS group. The lack of a specific structural collagen marker to describe the matrix associated with extrinsically aged skin is unsurprising due to the reduced collagen content favoring elastin fiber formation.

**Figure 4.**
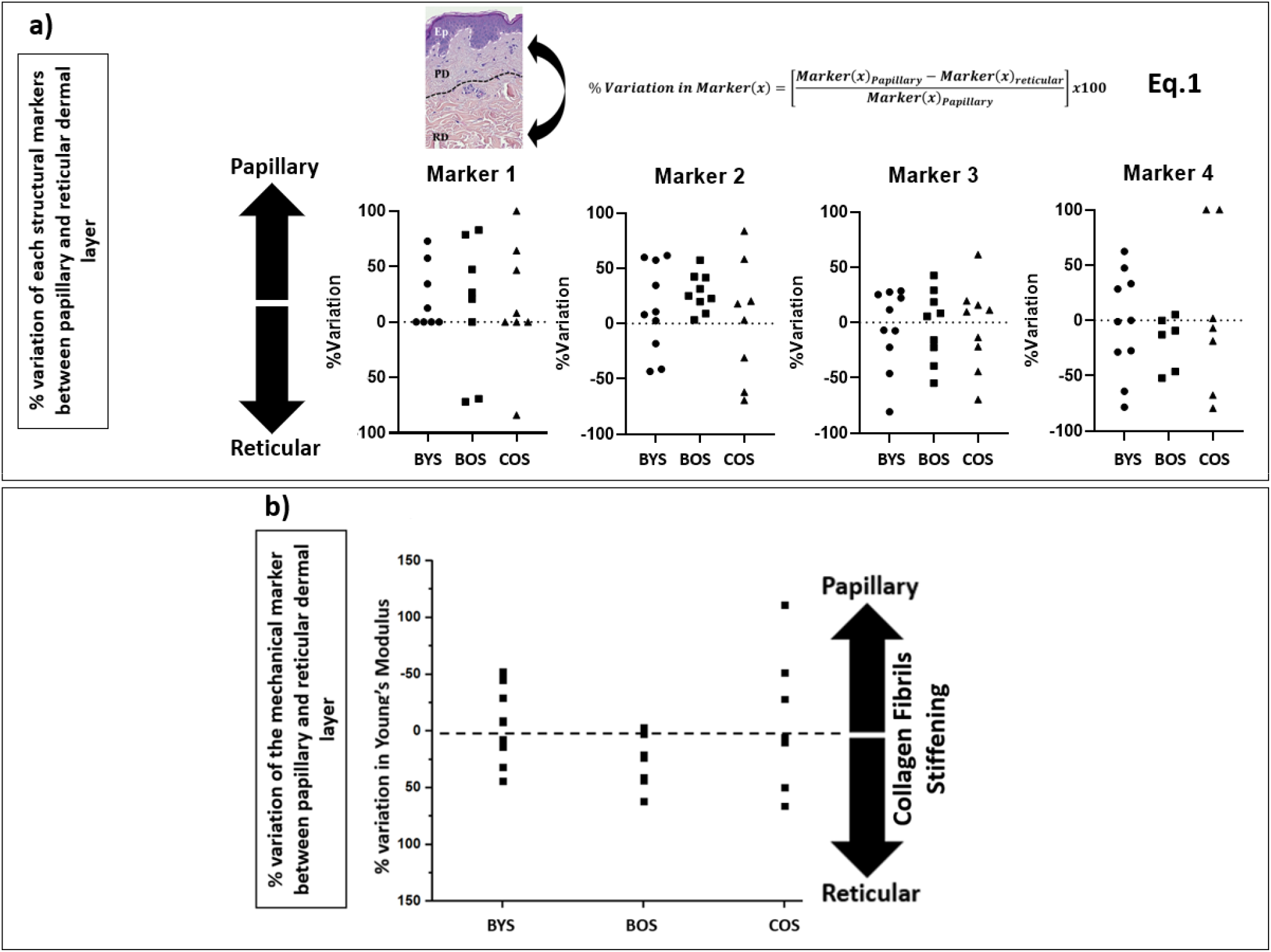
Evaluation of the prevalence of structural biomarkers for (type I) collagen as a function of dermal layers and skin groups: a) **%**variation in the prevalence of each nanoscale structural biomarkers for (type I) collagen between papillary and reticular dermis (defined in Eq. 1) as a function of the skin groups; b) variation in the prevalence of the mechanical biomarkers for (type I) collagen between papillary and reticular dermis (as defined in Eq. 1) for each skin group. The direction of the %variation indicates a greater marker prevalence in the papillary or reticular dermis.

### Collagen elasticity as a viable marker for age and anatomical differentiation

In a similar approach for the morphological assessment, we calculated the %variation of Young’s moduli between the papillary and reticular dermis, as presented in Figure 4b. This result shows that intrinsic aging promotes reticular collagen stiffening over papillary collagen within the intrinsically aged group (BYS compared to BOS). The extrinsically aged group (COS) did not show any trend toward stiffening of the papillary or reticular dermis. This heterogeneity in the mechanical response corroborates our finding in the nanohistology analysis, for which we could not use our pre-defined histomorphological markers. Numerous studies have explored the variations in the collagen mechanical properties at various scales as a function of aging and focused on calculating the nominal values for the elastic modulus of dermal collagen fibrils or scaffolds ^12,13,19,23,24^. The range of published elastic modulus values for dermal collagen fibrils varies across several orders ranging from a few 100 kPa ^24^ to a few GPa^13^. This wide range in the elastic modulus values is directly linked to the histological section conditioning, indentation size, inconsistent indentation load, limited sample size, and collagen hydration level. Unfortunately, it is challenging to compare study outcomes in terms of collagen elasticity measurement due to the wide variations in the techniques used.

### Discrimination between chronological and biological aging.

To explore from a multivariate perspective the correlation between the structural (marker 1-2-3-4) and mechanical (elasticity) markers regarding the chronological age of the donors, both unsupervised (regardless of the initially assigned group) PCA and supervised (with the age group as response variable) PLS-DA analyses have been carried out. Figure 5a presents the PCA loading plot associated with the %variation of the markers across all donors. The size of the vectors associated with the % variation in markers 3, 4, and elasticity confirm that these three markers are the more potent contributor to the principal components, thus, more prominent differentiators of the entire dataset. On the other hand, the size of the vector associated with marker 1 confirms marker 1 has a limited impact on the dataset. Therefore, we supplemented our PCA analysis with a PLS-DA using the donors’ age binary class as the response variable rather than the markers (Fig. 6b). The Biplot chart allows individuals and descriptors (% variation of all markers) to be simultaneously represented and allows us to interpret the individuals in terms of descriptors.

**Figure 5.**
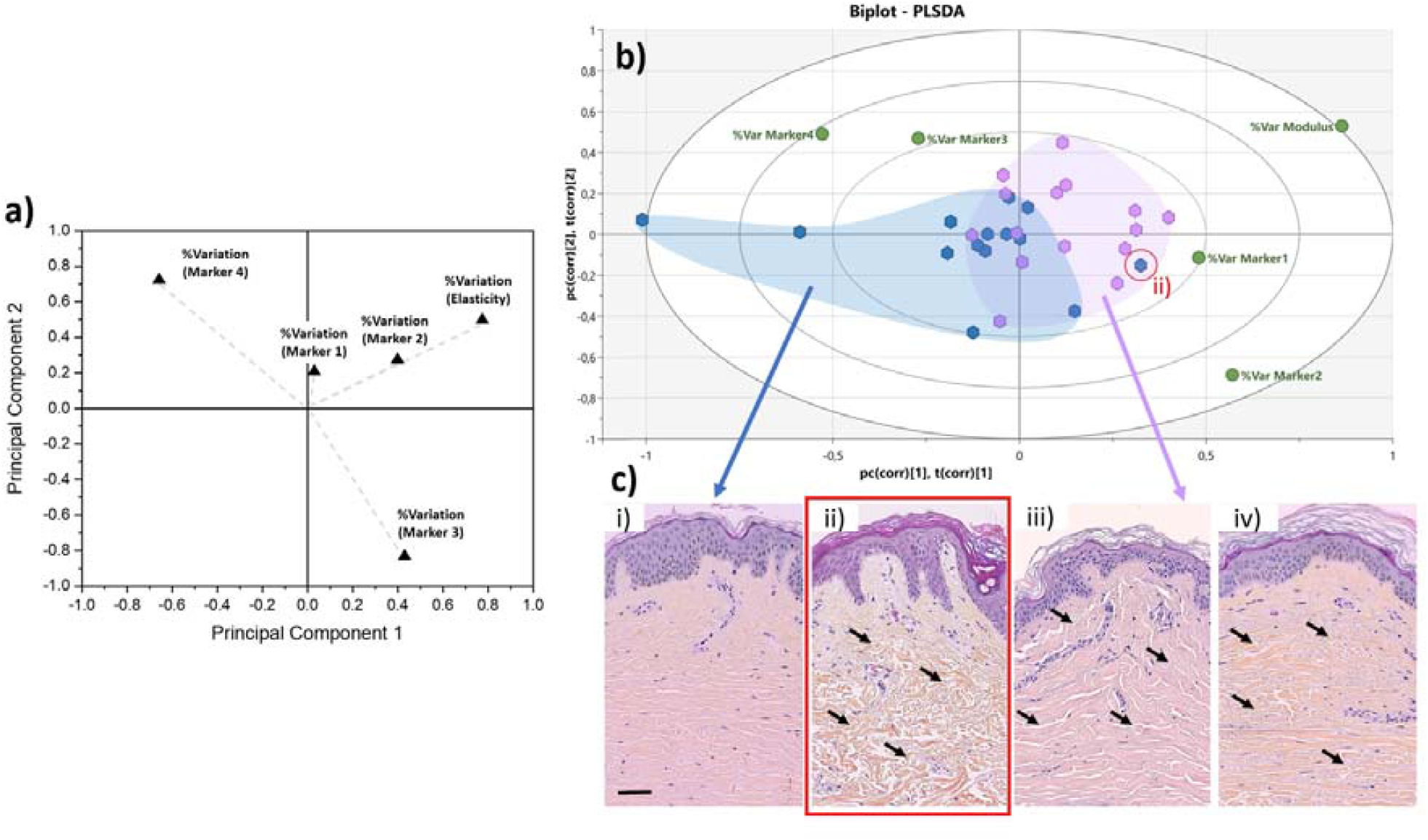
Discrimination between chronological and biological aging using multivariate analyses. a) 2-D PCA loading plot associated with the %variation of the markers across the entire dataset for all donors. Variables are presented as vectors extending away from the origin to assess their contribution towards the two principal components. b) Graphical biplot of the PLS-DA performed on the entire dataset using the donors’ age binary class as the response variable. The blue cluster regroups donors belonging to the young group (<30 years old - median 22.0±4.2 years old), whereas the purple cluster regroups donors belonging to the older group (>50 years old - median 66.0±10.4 years old). The red circled data point highlights an outlier belonging to the young group. representative HES Histology of Normal Human Skin from either the young or the older group, as presented in Figure 1. The red circle histological image refers to the outlier (young) case in b), which presents an old skin phenotype. The black arrows highlight the variations in the dermal layer morphology which can be described as atrophic and sparse in the case of ii), when compared to iii) and iv).

In this biplot, we can observe that the distribution of donors tends to form 2 clusters: all the young donors (<30y.o) are clustered on the left-hand side of the biplot (with two exceptions), whereas the older donors (>50 y.o) tend to cluster on the right-hand side of the biplot. Significant overlap exists between the two sub-populations around the central axis of the PLS-DA. This overlap can be associated with the well-known disparity between the chronological age of the donors and their biological age ^25^. Although a 20-year gap exists between our younger and old groups, one cannot assume that the chronological age of the donor is the same as their biological age. This is proven by the presence of a young donor case on the aged group’s outer edge (red circle). Figure 5c-ii presents the histological images of this young donor (22y.o). Despite the dermo-epidermal junction being non-uniform and presenting well-defined rete ridges, the underlying dermal collagen presents significant signs of aging-induced damage, as found in older donors (Figure 5c iii, iv). The dermal collagen is heavily atrophic and sparse for this young donor, likely associated with significant sun damage. The quantitative nanohistological assessment performed in this study could classify this donor’s collagen phenotype amongst the old group. This 22-year-old donor has the dermal fingerprint of a chronologically old donor, demonstrating the disparity between chronological and biological age again.

## SIGNIFICANCE

Exploring variations in the phenotypic properties of human largest and most accessible organs has been the subject of much research. However, quantifying the impact of chronic and pathological conditions on the structure and function of collagen at the sub-micron level remains challenging. This is due to a) a lack of current techniques to assess these properties systematically at the nanoscale and b) a lack of histopathology standards (at that scale) in the literature. By employing tools such as the Atomic Force Microscope, we can start cataloging the complexity of the dermal matrix at the nanoscale. All the studies to date present a snapshot of what the dermal matrix could be, but there is a significant lack of extensive studies. This poses a challenge for the dermo-cosmetic and dermo-pharmaceutical fields as our study demonstrates an extensive and diverse collagen phenotype across the donors’ ages with unique properties for everyone. This is especially true for the photo-exposed skin with the most heterogeneous biophysical properties.

On the contrary, the photo-protected skin remains more biomechanically homogeneous and presents some clear structural characteristics. Our study suggests that the relative difference in the collagen biophysical properties between the papillary and reticular dermis can be used as the most effective discriminator to differentiate between chronological and biological aging and the anatomical site. Furthermore, this differential phenotypic assessment of collagen at the nanoscale can be used to assess subtle variations in the collagen properties in dermal localized pathological conditions, such as Scleroderma ^19^ and skin cancer progression.

## CONCLUSION

Our study presents a significant milestone in understanding the complex nature of connective tissue, such as skin, as a function of aging. Here, we demonstrate that we can differentiate between individuals based on their biological age by considering individual skin samples as unique statistical entities with defined structural and mechanical variations in the reticular and papillary dermal collagen. Here, we used an AFM to image and quantify the collagen in the dermis before manually analyzing all the images. Using this approach supported by PCA, we demonstrated how the relative variations in structure and mechanics of the dermal collagen could be used to determine individuals’ biological age. Our approach requires advanced knowledge of collagen morphological variations at the nanoscale, for which no established standards exist. Yet, accessing the collagen structure at that scale is fundamental to help define candidate histological markers. As such, we are on course to create a new field of histology: quantitative nanohistology, to probe, quantify and explore connective tissues’ structural and functional properties at the sub-micron level.

## Acknowledgments

We would like to thank the London Centre for Nanotechnology (London UK) for granting access to the AFM Facility and UCL Consultants for their support.

